# Newly recruited intraepithelial Ly6A^+^CCR9^+^CD4^+^ T cells protect against enteric viral infection

**DOI:** 10.1101/2021.11.10.468106

**Authors:** Roham Parsa, Mariya London, Tiago Bruno Rezende de Castro, Bernardo Reis, Julian Buissant des Amorie, Jason G. Smith, Daniel Mucida

**Affiliations:** Laboratory of Mucosal Immunology, The Rockefeller University, New York, NY 10065, USA; Department of Microbiology, New York University Grossman School of Medicine, New York, NY 10016, USA; Laboratory of Lymphocyte Dynamics, The Rockefeller University, New York, NY 10065, USA; Molecular Cancer Research, Center for Molecular Medicine, University Medical Center Utrecht, Utrecht University, The Netherlands; Department of Microbiology, University of Washington School of Medicine, Seattle, WA 98109, USA; Howard Hughes Medical Institute, The Rockefeller University, New York, NY, USA

## Abstract

The intestinal epithelium comprises the body’s largest surface exposed to viruses. However, a role for intraepithelial T lymphocytes in resistance against viral infections remain elusive. By fate-mapping T cells recruited to the murine intestinal epithelium, we observed accumulation of CD4^+^ T cells after infection with murine norovirus (MNV) or mouse adenovirus type-2 (AdV), but not after reovirus infection. Intraepithelial CD4^+^ T cells recruited after MNV or AdV infection co-express Ly6A and CCR9, and exhibit T helper 1 and cytotoxic profiles. Although these cells display a diverse TCR repertoire, they conferred protection against AdV and MNV both *in vivo* and in an organoid co-culture model in an IFN-γ-dependent manner. Ablation of the T cell receptor (TCR) or the transcription factor ThPOK in CD4^+^ T cells prior to infection prevented viral control, while TCR ablation during infection did not impact viral clearance. These results uncover a protective role for intraepithelial Ly6A^+^CCR9^+^CD4^+^ T cells against enteric viruses.

## Introduction

The highly specialized intestinal immune system is charged with maintaining tolerance to harmless stimuli from commensal bacteria and food, while providing protective immunity against pathogens and epithelial cancers (Florsheim et al., 2021; Honda and Littman, 2016; McDonald et al., 2018; Tanoue et al., 2016). Intraepithelial lymphocytes (IELs) comprise a large T cell population located at the critical interface between the intestinal lumen and the core of the body. IELs can provide a first line of immunity in mice and humans, while balancing tolerance and defense (Bilate et al., 2020; Cervantes-Barragan et al., 2017; Cheroutre et al., 2011; Edelblum et al., 2015; Hoytema van Konijnenburg et al., 2017; Jabri and Sollid, 2009; Sujino et al., 2016). Among the IEL populations, CD4^+^ T cells are key players in intestinal homeostasis, finely tuning responses at the level of antigen recognition and functional differentiation. In the intestinal epithelium (IE) and underlying lamina propria (LP), tissue adapted pro-inflammatory, regulatory (Treg) and intraepithelial (CD8αα^+^ CD4_IEL_) CD4^+^ T cells coordinate immunity and tolerance to diverse intestinal stimuli (Cervantes-Barragan *et al.*, 2017; Honda and Littman, 2016; McDonald *et al.*, 2018; Sujino *et al.*, 2016). Both LP and IE CD4^+^ T cells are antigen-experienced and co-express markers of activation, gut-homing and tissue residency, including CD44, CD69, CD103 and CD8αα homodimers (Cepek et al., 1994; Kilshaw and Murant, 1990; Masopust and Soerens, 2019).

Enteric viruses, such as norovirus, rotavirus and adenovirus, are among the most common causes of acute gastroenteritis in humans and are responsible for morbidity and mortality worldwide (Iliev and Cadwell, 2021; Kapikian et al., 1972; Wilhelmi et al., 2003). These enteric viruses can infect intestinal epithelial cells and myeloid cells in the LP, provoking robust innate and adaptive immune responses (Iliev and Cadwell, 2021); however, a functional role for intestinal T cells in resistance to enteric viruses is not well established.

We sought to address the functional role of intestinal T cells in the response to enteric viral infections by using a novel T cell fate-mapping strategy to identify and characterize IE T cell dynamics during enteric viral infection. We observed specific recruitment and accumulation of CCR9^+^Ly6A^+^CD4^+^ T cells, expressing a T helper 1 (TH1) and cytotoxic profile, in the epithelium after murine norovirus (MNV) and mouse adenovirus-2 (AdV) infections. Virus-recruited CD4^+^ T cells displayed increased T cell receptor (TCR) diversity while ablation of the TCR or the CD4-lineage defining transcription factor T-Helper-Inducing POZ/Krueppel-Like Factor (ThPOK) in CD4^+^ T cells prior to AdV infection prevented viral control. In summary, we found that intraepithelial CD4^+^ T cells protect against enteric virus infection in a ThPOK- and IFNγ-dependent manner.

## Results

### Enteric virus infection leads to distinct T cell dynamics

To characterize T cell dynamics in the IE during enteric virus infections, we infected mice with four different viruses, chronic MNV (CR6), acute MNV (CW3) (Kernbauer et al., 2014), reovirus T1L (T1L) (Bouziat et al., 2017) and AdV (Wilson et al., 2017). CR6, CW3 and T1L infect myeloid and stromal cells in the LP (Baldridge et al., 2016; Mainou et al., 2013), and CR6 can also infect Tuft cells in the IE to establish long lasting chronic infection (Wilen et al., 2018). AdV exclusively infects and replicates in the IE (Hashimoto et al., 1970; Takeuchi and Hashimoto, 1976) (Figure S1A). We have previously shown that during steady state, CD4^+^ T cells enter the IE as CD103^-^ effector T cells and gradually acquire an IEL phenotype with progressive expression of CD103 and CD8αα (Bilate *et al.*, 2020; London et al., 2021). To understand T cell recruitment dynamics to the site of enteric infections, we first quantified the frequencies of CD103^-^ T cells in the IE 10 days post infection by flow cytometry. We observed that AdV infection resulted in an increase of CD4^+^CD103^-^ T cells whereas T1L infection led to an increase in CD8αβ^+^CD103^-^ T cells in the IE (Figure 1A, B). To independently assess peripheral T cell recruitment to the IE during viral infections, we utilized a novel fate-mapping approach. We crossed *Sell*^CreERT2^ (Merkenschlager et al., 2021) with *Rosa26*^CAG-LSL-tdTomato^ (iSell^Tomato^) to permanently label naïve T cells (CD62L^+^) as Tomato^+^ upon tamoxifen treatment, enabling a strategy that enriched for “ex-naïve” (CD62L^-^Tomato^+^) pathogen-specific T cells that migrated to the intestine (Figure 1C and Figure S1B). Analysis of tamoxifen-treated iSell^Tomato^ mice 10 days post infection revealed a significant increase of recently recruited (Tomato^+^) CD4^+^ T cells post AdV and CR6 infections, whereas T1L and CW3 infections led to a preferential recruitment of CD8αβ^+^ T cells (Figure 1D, E). Additionally, most enteric viruses led to a significant recruitment of both CD4^+^ and CD8αβ^+^ T cells to the LP (Figure S1C).

**Figure 1.**
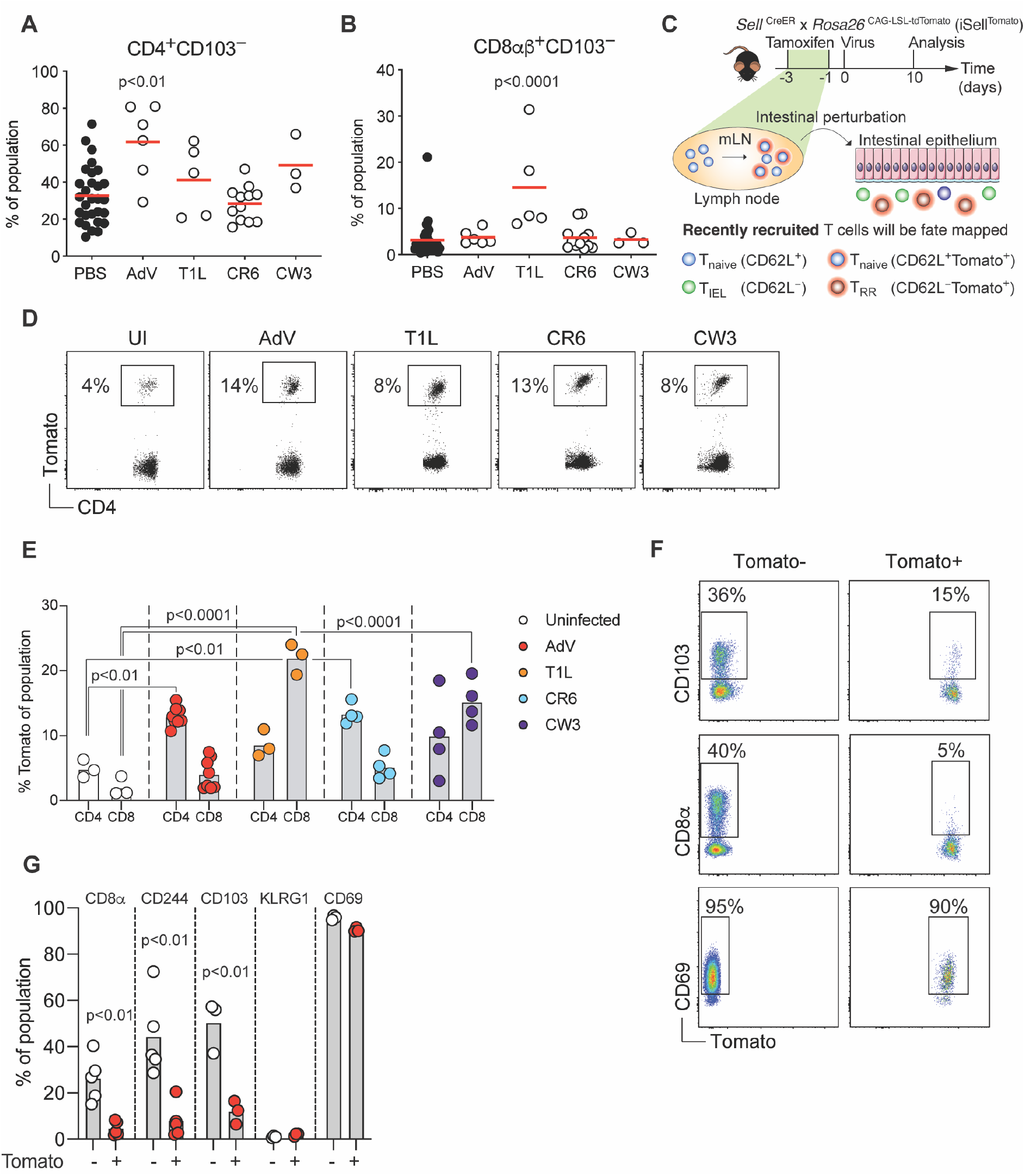
Distinct intraepithelial T cell dynamics post enteric viral infections. B6 mice were orally infected with 10^7^ infectious units (i.u.) of murine adenovirus-2 (AdV), 10^8^ plaque forming units (pfu) of reovirus T1L, 3×10^6^ pfu of either murine norovirus (MNV) CR6 or CW3. CD103^-^CD4^+^ (**A**) or CD103^-^CD8αβ^+^ (**B**) intraepithelial lymphocytes (IEL) were analyzed 10 days post infection among total CD4 or CD8αβ T cells, respectively. (**C**) Experimental overview of the T cell fate-mapping model. iSell^Tomato^ mice were treated orally with tamoxifen 1 and 3 days prior to viral infection, RR: recently recruited. (**D-E**) iSell^Tomato^ mice were orally infected with 10^7^ i.u. of AdV, 10^8^ pfu of T1L, or 3×10^6^ pfu of either CR6 or CW3, TCRβ^+^CD4^+^CD62L^-^ and TCRβ^+^CD8αβ^+^CD62L^-^ IELs were analyzed for tomato expression 10 days post infection. (**D**) Representative dot plots of tomato expression among CD4^+^ IELs. (**E**) Frequencies of tomato expression among CD4^+^ or CD8^+^ IELs. (**F-G**) iSell^Tomato^ mice were infected with 10^7^ i.u. of AdV and expression of CD103, CD8α and CD69 were analyzed among TCRβ^+^CD4^+^CD62L^-^Tomato^+^ or Tomato^-^ cells (**F**) Representative surface expression of markers as indicated among Tomato^+^ and Tomato^-^ T cells. (**G**) Frequencies of Tomato+ or Tomato-cells expressing the indicated markers. Data are expressed as means of individual mice (n = 3–5 of two independent experiments in D-G). p values are as indicated, one-way ANOVA plus Bonferroni test in A and E, Student’s t-test in G.

We then focused on AdV infection as this virus preferentially recruited CD4^+^ T cells and exclusively infects the intestinal epithelium. As expected based on a prior study, newly recruited CD4^+^ T cells in response to AdV infection expressed low levels of typical IEL markers such as CD103, CD8αα or CD244 (Reis et al., 2013), but the majority of Tomato^+^ T cells expressed CD69 indicating recent activation, or early tissue-residency (Faria et al., 2017) (Figure 1F, G and Figure S1D). Using a novel fate-mapping strategy we revealed that the dynamics of recently recruited T cells differ between enteric viral infections.

### Recruited intraepithelial CD4^+^ T cells display a mixed T_H_1 and T cytotoxic profile

Next, we analyzed the gene expression profiles of newly recruited IE and LP CD4^+^ T cells from AdV-infected mice. Unbiased clustering of the gene expression data and principal component analysis segregated CD4^+^Tomato^+^ T cells from infected mice in comparison to control mice (Figure 2A, B). CD4^+^Tomato^+^ T cells from AdV infected mice displayed tissue specific gene expression such as *Ccl5* and *Itga1* in the IE, and *Fasl* and *IL12rb2* in the LP, indicating tissue adaptation and some degree of compartmentalization (London *et al.*, 2021). CD4^+^Tomato^+^ T cells from both compartments of AdV infected mice displayed expression of inflammation- and gut residency-associated genes (*Ccr9, Ly6a, Lztfl1* and *Fyco1*), with a mixed T_H_1 (*Prdm1, Tbx21, Id2* and *Cxcr6)* and T cytotoxic (CTL) (*Runx3, Nkg7, Gzma* and *Gzmk*) profiles (Figure 2A, C and D). Overall, these results suggest that intestinal CD4^+^ T cells recruited upon AdV infection display mixed T_H_1 and CTL profiles.

**Figure 2.**
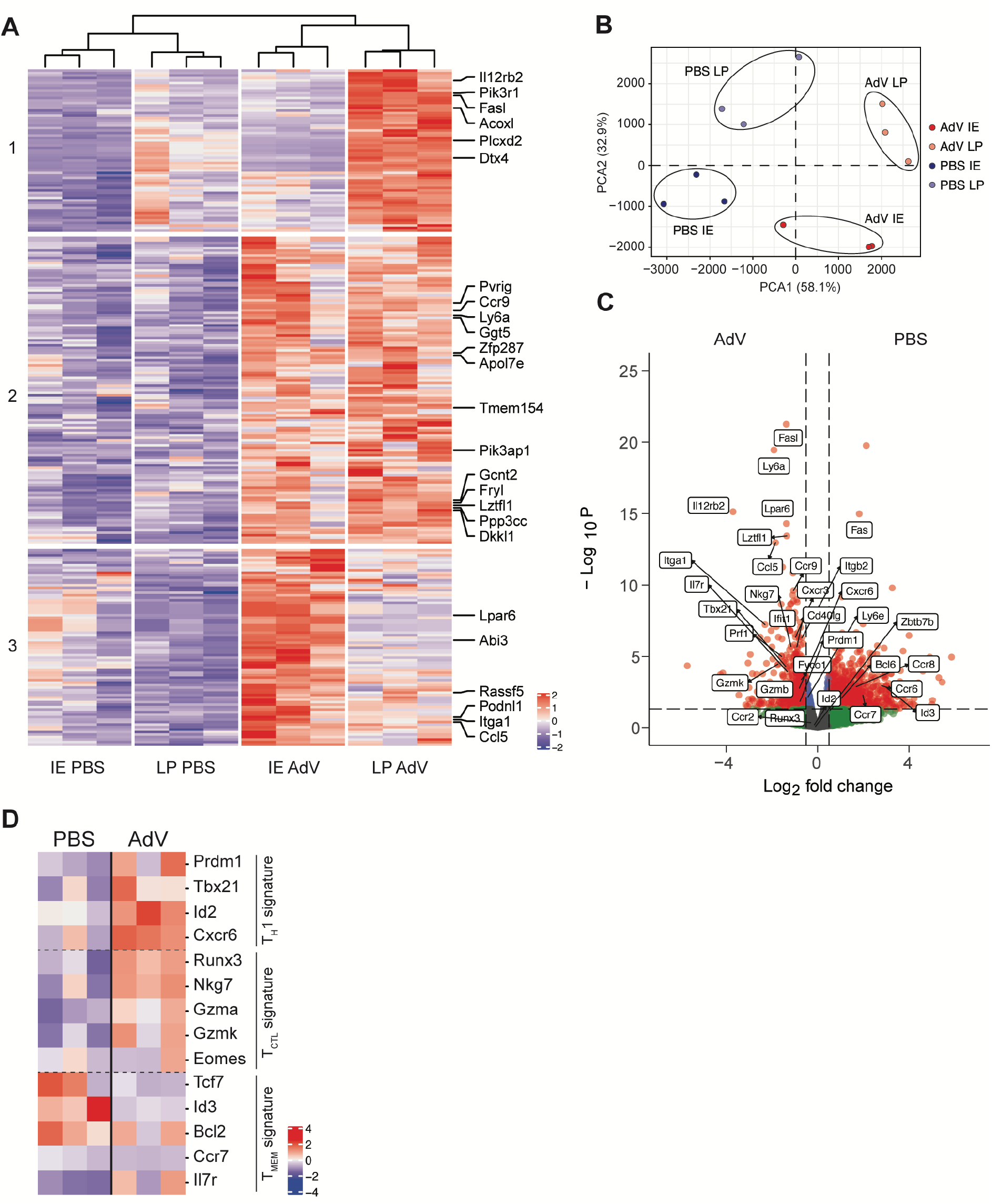
Transcriptional analysis of epithelial-recruited CD4^+^ T cells post AdV-infection. iSell^Tomato^ mice were infected with AdV or treated with PBS vehicle control, and CD4^+^CD62L^-^Tomato^+^ T cells were sorted from lamina propria (LP) or intestinal epithelium (IE) 10 days post infection for bulk RNAseq (**A**) Heatmap clustering of the most significant differentially expressed genes represented by normalized Z score in LP and IE of AdV-infected and PBS treated control mice. Top 25 genes are annotated. FDR < 0.05 (**B**) Principal component analysis of sorted T cells in IE and LP compartment of AdV-infected and control mice (**C**) Volcano plot visualizing gene fold change (X-axis) versus p value (Y-axis) between T cells from AdV-infected and control mice for both IE and LP combined. Dashed line on y-axis denotes p value of 0.05 and dashed line on x-axis denotes fold change of ± 2. (**D**) Transcriptional signatures represented by normalized Z score of T cells between AdV-infected and control mice for both IE and LP combined. n = 3 mice per group. Data are expressed as means of individual mice.

Our transcriptional analysis indicated a heterogenous profile of the recruited CD4^+^ IE T cells post enteric AdV infection. To initially assess their functional profile, we infected tamoxifen-treated iSell^Tomato^ mice and characterized CD4^+^Tomato^+^ T cells 10 days post AdV infection by using markers such as Ly6A and CCR9, enriched in our RNAseq data, using flow cytometry. As expected, we detected only a few fate-mapped CCR9^+^Ly6A^+^CD4^+^ T cells in the IE and LP compartments in naïve uninfected mice. In contrast, AdV-infection significantly increased the recruitment of CCR9^+^Ly6A^+^CD4^+^ T cells and granzyme B (GzmB) expression (Figure 3A). Furthermore, the majority of recruited CD4^+^Tomato^+^ in AdV-infected mice expressed the transcription factor *Tbx21* (T-bet) (Figure 3B), associated with T_H_1 phenotype as well as with IEL differentiation (Reis et al., 2014). Accordingly, we detected increased IFN-γ expression in CD4^+^Tomato^+^ IE T cells from AdV-infected mice in comparison to CD4^+^Tomato^+^ IE T cells isolated from uninfected mice (Figure 3C).

**Figure 3.**
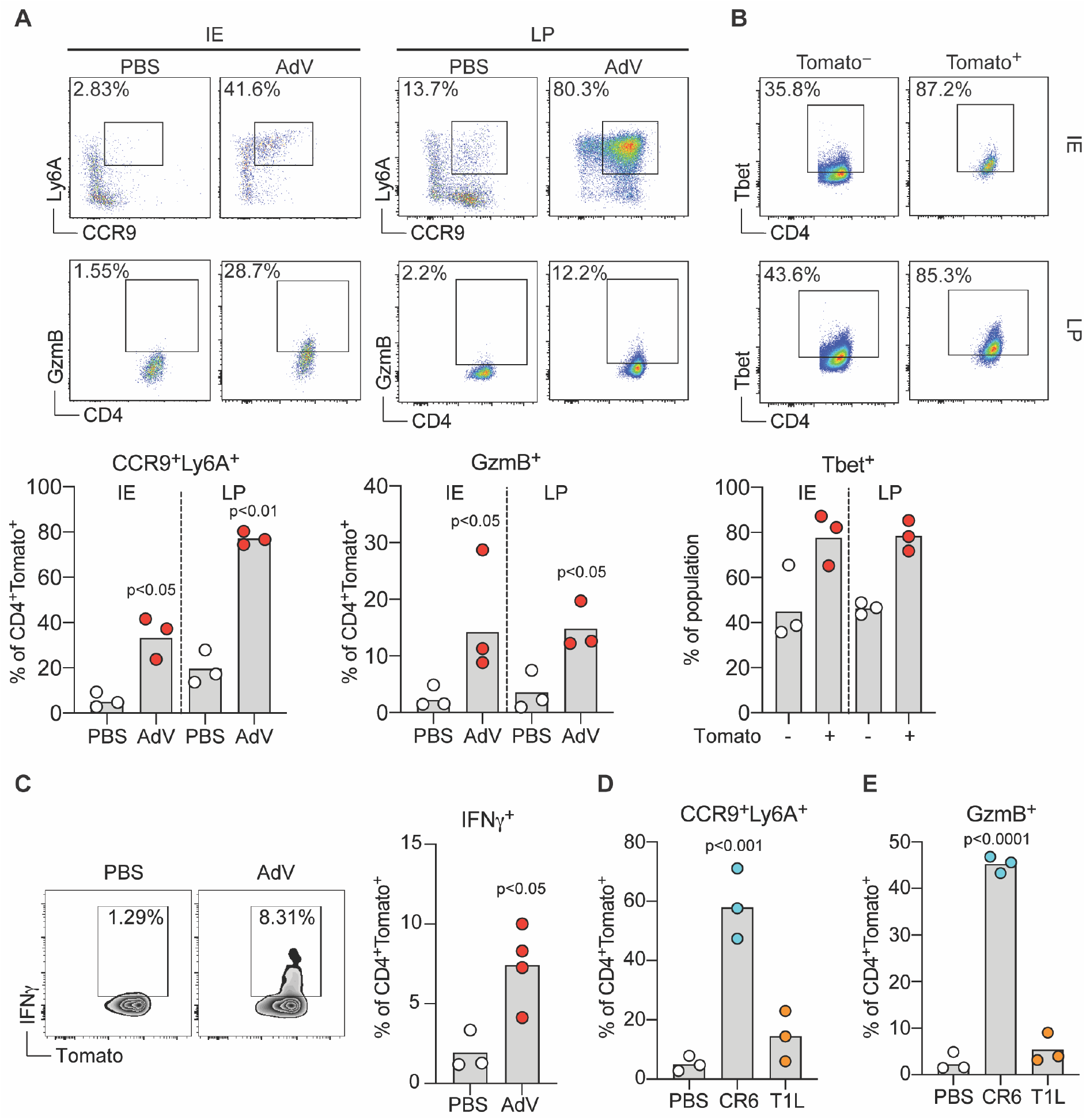
AdV- and CR6-recruited IE CD4^+^ T cells are enriched for CCR9^+^Ly6A^+^ cells and acquire a T_H_1 and cytotoxic profile. iSell^Tomato^ mice were infected with AdV or treated with PBS vehicle only, and CD4^+^CD62L^-^Tomato^+^ T cells were analyzed 10 days post infection. (**AC**) Representative plots (top) of marker expression and population frequencies (bottom) as indicated among CD4^+^CD62L^-^Tomato^+^ T cells in LP or IE compartments as indicated (**A**). (**B**) Representative plots (top) and frequencies (bottom) of T-bet expression among Tomato^+^ or Tomato^-^ cells in IE. (**C**) Representative plots (left) and frequencies (right) of IFN-γ production among CD4^+^CD62L^-^Tomato^+^ T cells in the IE. (**D-E**) iSell^Tomato^ mice were infected with CR6 or T1L, or treated with PBS vehicle only, and CD4^+^CD62L^-^Tomato^+^ T cells was analyzed for Ly6A, CCR9 (**D**) and GzmB (**E**) expression 10 days post infection among CD4^+^CD62L^-^Tomato^+^ cells. Data are expressed as means of individual mice (n = 3–5 of two independent experiments). p values are as indicated, Student’s t-test in A and C. One-way ANOVA plus Bonferroni test in D and E.

Because we observed that different enteric viruses give rise to distinct T cell responses in the IE compartment (see Figure 1E above), we infected tamoxifen-treated iSell^Tomato^ mice with CR6 or T1L to determine if these viruses could also induce CCR9^+^Ly6A^+^CD4^+^ T cells. In agreement with differences observed in the CD4 T cell recruitment between these viruses, CR6, but not T1L infection, led to increased CCR9^+^Ly6A^+^CD4^+^ T cells (Figure 3D) with similar GzmB expression as observed in AdV-recruited CD4^+^Tomato^+^ T cells (Figure 3E). These results reveal that MNV and AdV infections induce epithelial recruitment of CD4^+^ T cells that express the gut-homing receptor CCR9, the activation marker Ly6A, and an overall T_H_1/CTL phenotype.

### Recruited intraepithelial CD4^+^ T cells are clonally diverse and temporally TCR-dependent for controlling AdV-replication *in vivo*

To determine a role of the T cell receptor (TCR) in controlling AdV viral replication, we first extracted the TCR CDR3 sequences for both the a and the β chain from our bulk RNAseq data. Analysis of the CDR3 sequences suggested an increased number of reads for unique CDR3 sequences in CD4^+^ T cells recruited during acute viral infection in comparison to CD4^+^ T cells recently recruited in naïve mice (Figure 4A). We have recently shown that CD4^+^ T cells in the IE undergo clonal selection and reduction in TCR diversity as cells differentiate from conventional CD103^-^ CD4^+^ T cells into differentiated CD103^+^CD8αα^+^ CD4^+^IELs (Bilate *et al.*, 2020).

**Figure 4.**
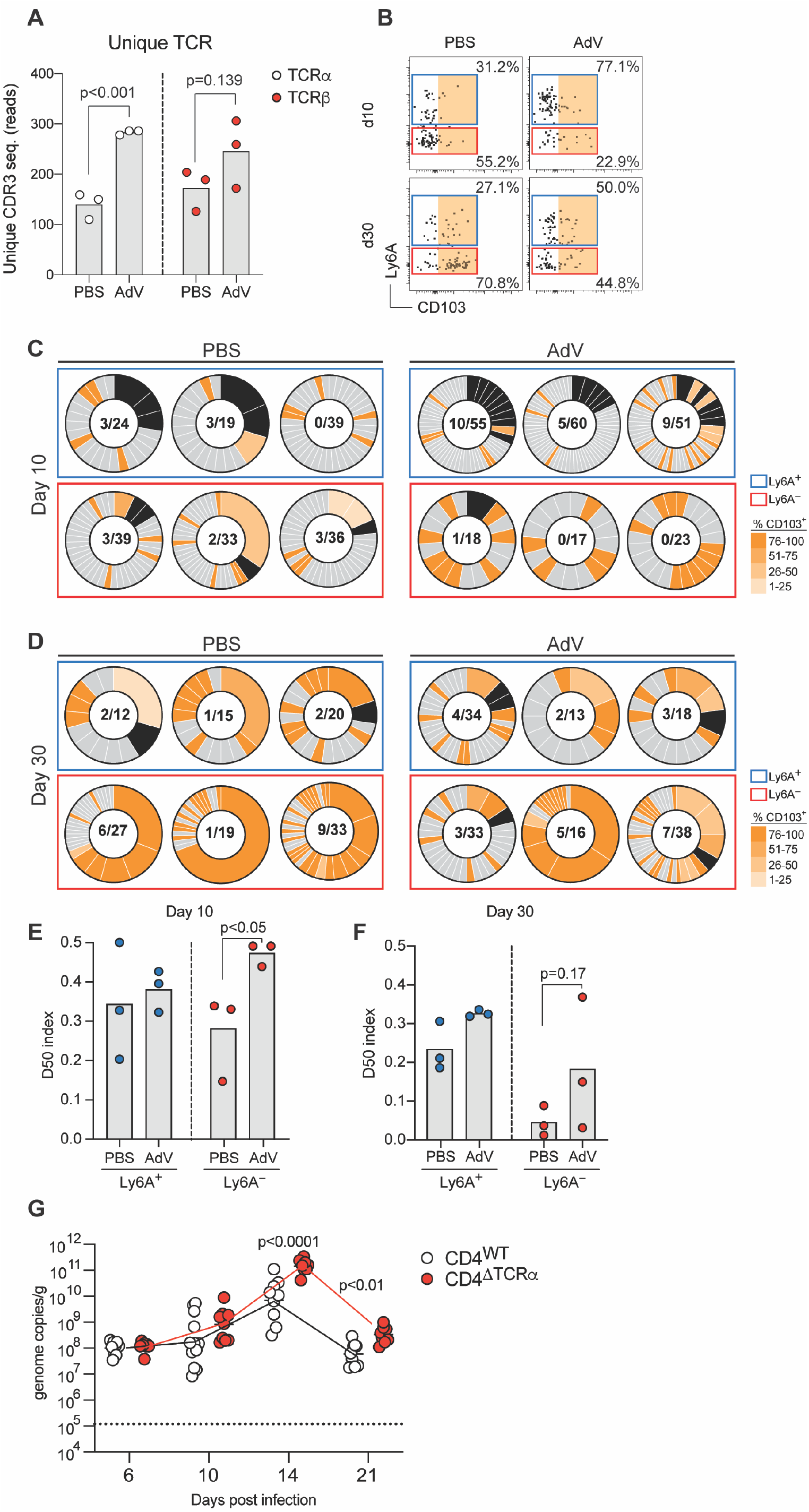
Clonally diverse Ly6A^+^CD4^+^ T cells are recruited to the IE compartment 10- and 30-days post AdV-infection. (**A**) Extraction of unique TCRα and TCRβ CDR3 sequence reads from bulk RNAseq data in figure 2. **(B-F)** iSell^Tomato^ mice were infected with AdV or treated with PBS vehicle control only, and CD4^+^CD62L^-^Tomato^+^ T cells were single-cell sorted at 10- and 30-days post infection for TCRβ-seq. (**B**) Ly6A and CD103 expression analysis of CD4^+^CD62L^+^Tomato^+^ T cells indexed and sorted for analysis in B and C. Colored gates represent indexed analysis in B and C. Orange area indicates CD103^+^ gate for C and D. Single cell TCRβ analysis at day 10 (**C**) and day 30 (**D**) post AdV-infection. Box colors represent gate color in A. Black pie regions represent expanded clones, grade of orange represents level of CD103 expression per clone indexed in A. Nominator represents total number of expanded clones, denominator represent total number of sequenced clones. (**E-F**) D50 analysis of CD4^+^Tomato^+^Ly6A^+^ and Ly6A^-^ at day 10 (**E**) and day 30 (**F**) post AdV infection. (**G**) Viral genome copies of AdV per gram of feces of AdV-infection over time of iCD4^ΔTCRα^ mice, dashed line represents limit of detection. Data are expressed as mean of individual mice and n = 3 mice per group for A-F, n = 5–12 (minimum two independent experiments in G). Significant p values as indicated, Student’s t-test in A, E, F and G.

To further investigate the TCR repertoire of AdV-recruited IE CD4^+^ T cells, we infected tamoxifen-treated iSell^Tomato^ mice with AdV and sequenced the TCRβ chain of single cell sorted Tomato^+^ CD4^+^ T cells, 10- and 30-days post-infection (AdV) or post-PBS injection (PBS). As shown earlier, most of the Tomato^+^ CD4^+^ T cells from AdV-infected mice were Ly6A^+^, whereas Tomato^+^ CD4^+^ T cells in uninfected mice were mostly Ly6A^-^ (Figure 4B).

Analysis of newly recruited (Tomato^+^) Ly6A^-^ CD4^+^ T cells from uninfected mice also confirmed our previous observations, revealing clonal expansions with reduced TCR diversity and increase in CD103 expression in most of the clones at day 30 [Figure 4B-F; TCR diversity was assessed by the diversity 50 index (D50), indexes varied from least diverse; 0, to most diverse; 0.5]. However, contrary to expectations for T cell responses to pathogens, newly recruited (Tomato^+^) IE CD4^+^ T cells from AdV-infected mice, regardless of Ly6A expression, did not show reduced TCR diversity and higher clonal expansions. Instead, CD4^+^ T cells from AdV-infected mice displayed similar or even less clonal expansion than control mice and the Ly6A^-^ population showed significantly higher diversity at day 10 post-infection (Figure 4C-F). Furthermore, recruited CD4^+^Tomato^+^ T cells in AdV-infected mice displayed limited IEL differentiation, as assessed by CD8αα and CD103 expression at day 30 post-infection, with and remained similarly diverse (Figure 4B and Figure S2A).

To initially investigate possible anti-viral functions of gut recruited CD4^+^ T cells, we first assessed a role for TCR expression by these cells. In a recent study, we found that intraepithelial CD4^+^ T cells require TCR expression during their differentiation from peripheral precursors, but not necessarily for maintenance of their epithelial program, or function in a model of bacterial infection (Bilate *et al.*, 2020). We infected mice in which we inducibly deleted TCRα in CD4^+^ T cells (*Cd4^CreER^*x*Trac*^fl/fl^, iCD4^ΔTCRα^) immediately before AdV-infection. We found increased AdV titers in the stool of iCD4^ΔTCRα^ mice, pointing to a role for the TCR in CD4^+^ T cell control of viral replication in the intestine (Figure 4G). However, analogous to findings obtained in our previous study (Bilate *et al.*, 2020), excising TCRα from CD4^+^ T cells 6 days post infection did not impact anti-viral mechanisms during primary, or memory responses (Figure S2B, C). Hence, recruited IE CD4^+^ T cells post AdV infection retained clonal diversity and the activation marker Ly6A with limited expression of CD103. In contrast, recruited CD4^+^ T cells in uninfected mice, had reduced clonal diversity and increased CD103 expression over time. Furthermore, our data suggest for a requirement of the TCR during CD4^+^ T cell activation but removal of the TCR post infection did not impact anti-viral mechanisms.

### AdV- and CR6-recruited IE CD4^+^ T cells clear viruses in intestinal organoid cultures

To further examine the ability of virus-recruited CD4^+^ T cells to regulate viral replication, we developed an intestinal organoid (Sato et al., 2009) and T cell co-culture system *in vitro.* To visualize ongoing AdV infection and replication in the organoids, we used a replication competent mouse adenovirus-2 encoding green fluorescent protein (GFP) fused to the minor capsid protein XI (AdV-GFP) (Wilson *et al.*, 2017). AdV-GFP-infected organoids displayed GFP signal as early as 6 hours post-infection; when cultures were continuously monitored for up to 48 hours (Figure 5A, B and Suppl. Movie 1). We sorted CD4^+^Tomato^+^ IE T cells isolated from tamoxifen-treated iSell^Tomato^ mice infected with AdV, CR6 or T1L, 10 days post infection and co-cultured them with AdV-GFP infected organoids (Figure 5C). CD4^+^Tomato^+^ IE T cells isolated from either AdV- or CR6-infected mice, but not cells isolated from T1 L-infected mice, controlled AdV-GFP replication in intestinal organoids, suggesting that the differentiation of CCR9^+^Ly6A^+^CD4^+^ T cells is associated with cross-protection against enteric viruses (Figure 5D and E), data that indirectly reinforces the observations that continuous TCR expression by virus-recruited CD4^+^ T cells may not be required to regulate viral replication. To specifically address a functional role by recruited Tomato^+^CD4^+^ T cells in controlling AdV replication, we sorted Tomato^+^CCR9^+^Ly6A^+^, Tomato^+^CCR9^-^Ly6A^-^, Tomato^-^CCR9^+^Ly6A^+^ and Tomato^-^CCR9^-^Ly6A^-^ IE CD4^+^ T cells from tamoxifen-treated iSell^Tomato^ mice infected with AdV 10 days post infection and co-cultured them with AdV-GFP infected organoids. We found that Tomato^+^CCR9^+^Ly6A^+^CD4^+^ T cells had significantly higher capacity for controlling viral replication in the organoids in comparison to other T cell populations (Figure 5F).

**Figure 5.**
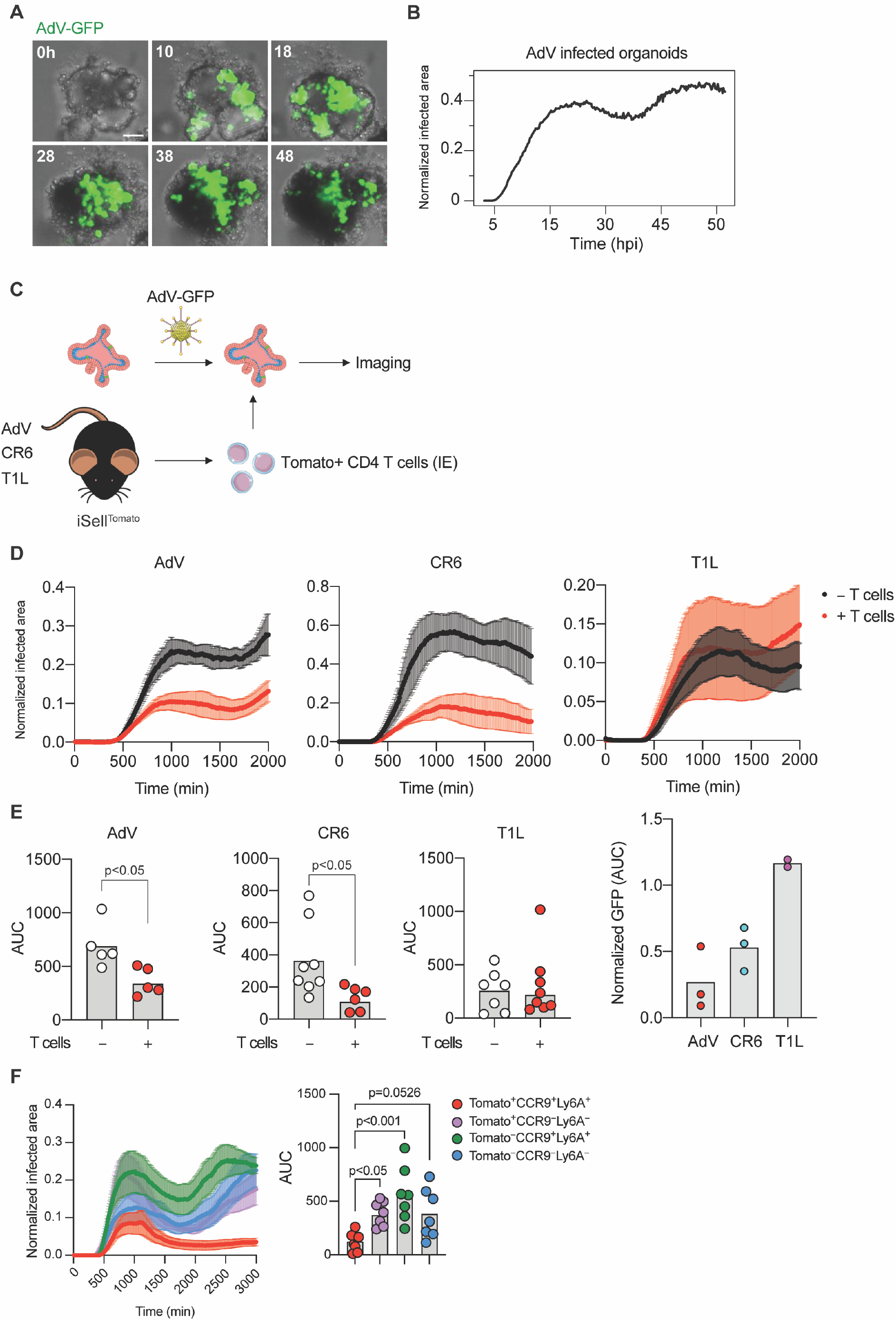
Newly recruited IE CD4^+^ T cells show anti-viral activity in AdV-infected intestinal organoid co-cultures. (**A-B**) Small intestinal organoids were infected with 10^4^ i.u. of AdV-GFP and imaged for 48 hours. (**A**) Representative images of organoids infected with AdV-GFP (green) at the indicated times post infection (hours, top left corner). (**B**) GFP expression area over time normalized to the area of organoids. (**C-E**) iSell^Tomato^ mice were infected with different viruses, CD4^+^Tomato^+^ IE T cells were then sorted and co-cultured with 10^4^ i.u. AdV-GFP infected organoids. (**C**) Experimental overview. (**D**) Representative graphs of GFP expression area from AdV-GFP infected organoids normalized to the area of the organoids over time with (red) or without (black) CD4^+^Tomato^+^ IE T cells derived from mice infected with AdV, CR6 or T1L 10 days post infection. Data are expressed as mean ± SEM of individual experiment (**E**) Summarized area under the curve (AUC) for D. Data points for individual organoids in E (left). Pooled data from 2-3 experiments (right). (**F**) iSell^Tomato^ mice were infected with AdV, CD4^+^Tomato^+^CCR9^+^Ly6A^+^, CD4^+^Tomato^+^CCR9^-^Ly6A^-^, CD4^+^Tomato^-^ CCR9^+^Ly6A^+^ and CD4^+^Tomato^-^CCR9^-^Ly6A^-^ IE T cells were then sorted and co-cultured with 10^4^ i.u. AdV-GFP infected organoids. Data are expressed as mean of individual organoid in E (except right figure of pooled data) and F. n = 5–10 organoids of minimum two independent experiments A-E, one experiment in F. p values as indicated, Student’s t-test in E or one-way ANOVA plus Bonferroni test in F.

We have previously correlated changes in T cell motility with anti-pathogen responses by IELs and detected preferential displacement of IELs to regions containing pathogens using live intravital microscopy (Hoytema van Konijnenburg *et al.*, 2017). Longitudinal imaging analysis of the organoid and T cell co-cultures showed cellular interactions between CD4^+^ IE T cells derived from AdV infected mice with AdV-infected epithelial cells (Figure 6A).

**Figure 6.**
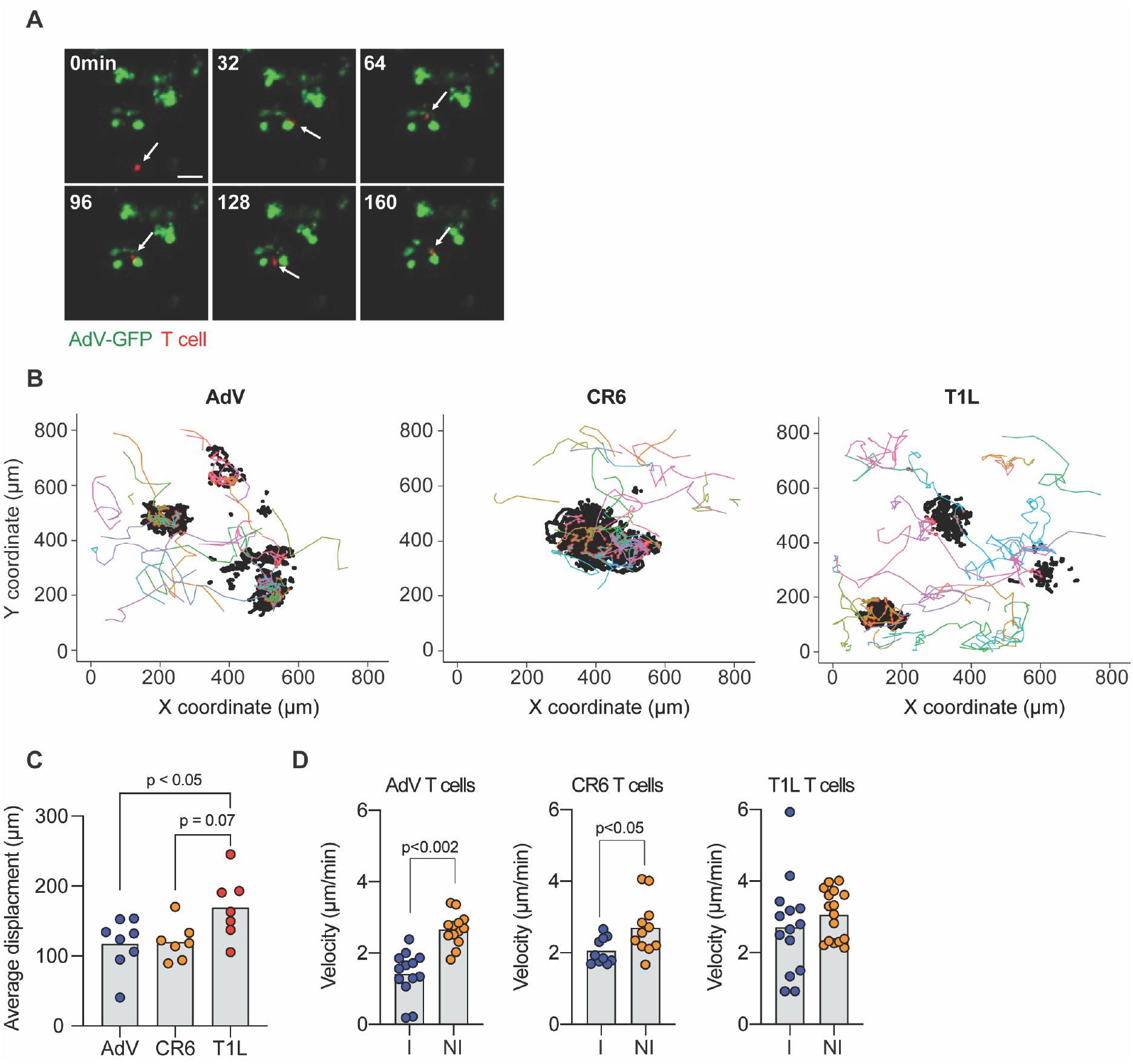
Newly recruited IE CD4^+^ T cells interact with AdV-infected epithelial cells in intestinal organoids. CD4^+^Tomato^+^ IE T cells derived from AdV, CR6 or T1L infected mice were co-cultured with AdV-GFP infected organoids and tracked over 48 ± 2 hours. (**A**) Representative images of AdV-GFP infected organoids (green) and IE CD4^+^Tomato^+^ T cells (red) derived from AdV-infected mice. Arrows indicate tracked T cell, top left corner indicates time post infection in minutes. (**B**) Tracking was obtained in 3 dimensions (x, y, and z); figure shows only x and y. Each colored line represents a unique T cell track, black dots represent GFP^+^ epithelial cells and red dots represent T cell-organoid interaction points (<20 μm radius distance). (C) Average displacement distance of tracked Tomato^+^ T cells co-cultured with AdV-infected organoids (**D**) Velocity of tracked tomato^+^ T cells, interacting T cells (I) within 20 μm radius of GFP^+^ cells, or non-interacting (NI) T cells outside the 20 μm radius of GFP^+^ cells. Data are expressed as average of pooled T cell tracks (n = 5–10 of minimum two independent experiments). p values as indicated, One-way ANOVA plus Bonferroni test in C, Student’s t-test in D.

Tracking of CD4^+^Tomato^+^ IE T cells from AdV- or CR6-infected mice suggested preferential migration of T cells towards the organoids, which contrasted with CD4^+^ T cells isolated from T1L-infected mice (Figure 6B, C and Suppl. Movies 2-4). Moreover, quantifying cellular velocity revealed that CD4^+^Tomato^+^ T cells, derived from AdV- or CR6-infected mice, interacting (<20 μm distance) with GFP^+^ epithelial cells slowed down in comparison to noninteracting T cells (>20 μm distance) from the same mice, an occurrence not observed in CD4^+^Tomato^+^ IE T cells derived from T1L-infected mice (Figure 6D). Therefore, the observed cross-protection by CD4^+^Tomato^+^ IE T cells derived from AdV- or CR6-infected mice correlates with distinct behavior and proximity to AdV-infected epithelial cells.

### ThPOK-dependent CCR9^+^Ly6A^+^CD4^+^ IE T cells cross-protect against enteric viruses

Upon arrival to the intestinal epithelium, peripheral CD4^+^ T cells gradually lose the helper T cell lineage-defining transcription factor ThPOK while upregulating the CD8 T cell lineage-defining transcription factor Runt-Related Transcription Factor 3 (Runx3), and T-bet (London *et al.*, 2021; Mucida et al., 2013; Reis *et al.*, 2013; Setoguchi et al., 2008; Sujino *et al.*, 2016; Taniuchi et al., 2002). This transition is accompanied by the loss of hallmarks associated with T helper subsets, including Rorγt or FoxP3 expression, and acquisition of IEL markers including CD244 and CD8αα homodimers (Mucida *et al.*, 2013; Reis *et al.*, 2013). Of note, while we had observed upregulation of granzyme B and IFN-γ by recruited CD4^+^ T cells during viral infection, which are characteristics of IEL differentiation, these cells do not display ThPOK downregulation or express CD8αα (see Figure 1G and Figure S2A above). To determine the role of transcription factors associated with IEL differentiation or CD4 T cell identity in the function and differentiation of gut-recruited CD4 T cells during enteric virus infection, we inducibly and specifically targeted *Zbtb7b* (iCD4^ΔThPOK^), *Tbx21* (iCD4^ΔTbet^) or *Runx3* (iCD4^ΔRunx3^) in CD4^+^ T cells upon tamoxifen administration prior to infection. AdV-infected iCD4^ΔThPOK^ mice displayed significantly reduced viral clearance compared to wild-type (WT) infected controls (Figure 7A), while iCD4^ΔRunx3^ mice presented delayed clearance but similar control by day 21 post-infection (Figure 7C). iCD4^ΔTbet^ showed no differences, compared to WT controls (Figure 7B). Consistent with a functional role for CCR9^+^Ly6A^+^CD4^+^ T cells in viral control, AdV-infected iCD4^ΔThPOK^ mice did not show recruitment of these cells to the IE compartment 10 days post infection, while iCD4^ΔTbet^ and iCD4^ΔRunx3^ mice displayed significant accumulation similar to control mice (Figure 7D). Furthermore, CCR9^+^Ly6A^+^CD4^+^ T cells from AdV-infected iCD4^ΔRunx3^ and iCD4^ΔThPOK^ mice showed reduced IFN-γ production when compared to infected control or iCD4^ΔTbet^ mice (Figure 7E), suggesting that unlike typical T_H_1 cells, CCR9^+^Ly6A^+^CD4^+^ T cell IFN-γ production is Tbet-independent.

**Figure 7.**
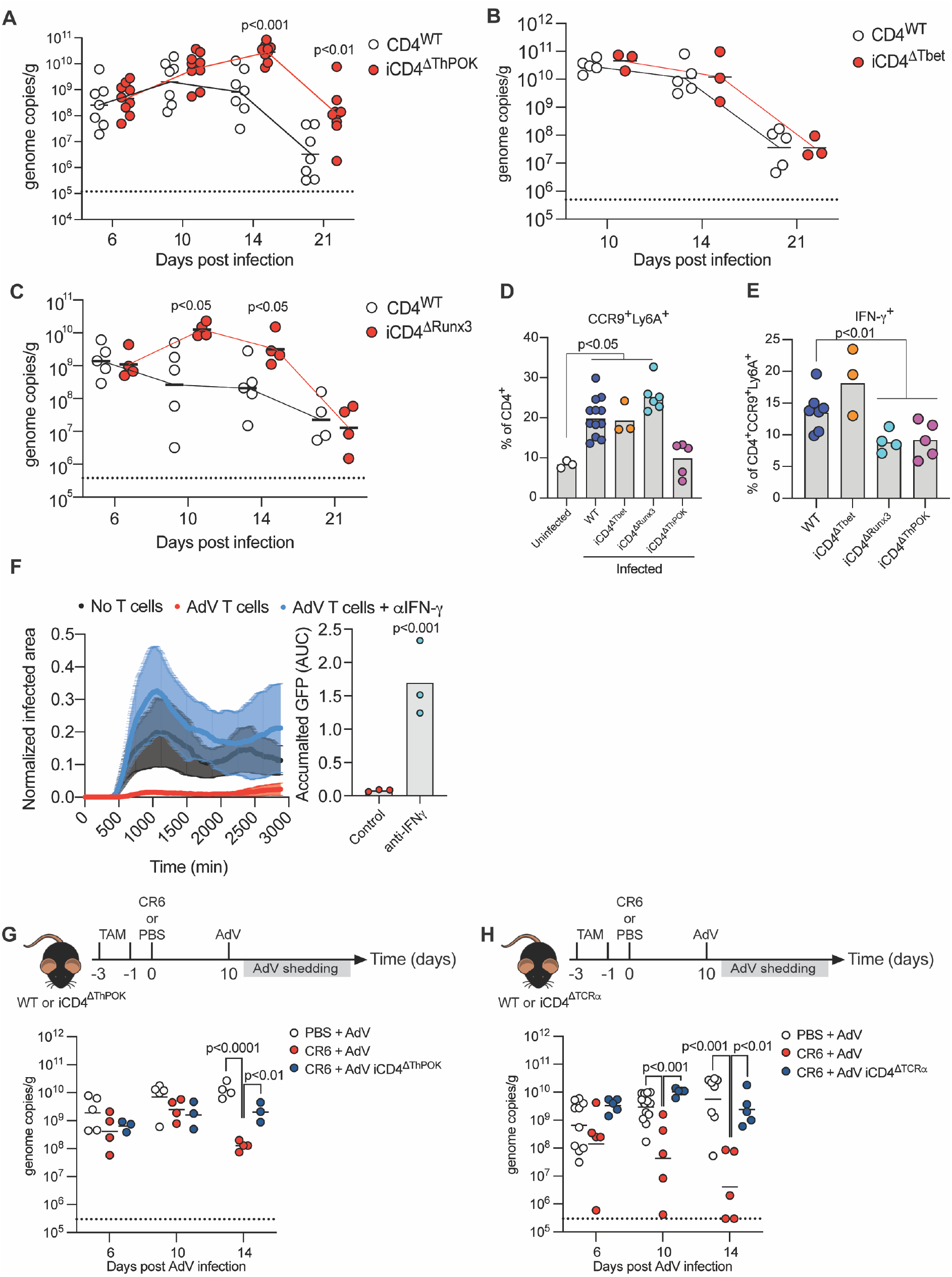
ThPOK-dependent CCR9^+^Ly6A^+^ CD4^+^ T cells show anti-viral functions *in vivo.* **(A-C)** AdV genome copies per gram of feces over time of post-infection (**A**) iCD4^ΔThPOK^, (**B**) iCD4^ΔTbet^ or (**C**) iCD4^ΔRunx3^ mice. Dashed line represents limit of detection. (**D-F**) Mice were infected with AdV and IE T cells were analyzed 10 days post infection. (**D**) Frequency of CCR9^+^Ly6A^+^ cells among CD4^+^ T cells. (**E**) IFN-γ production among CCR9^+^Ly6A^+^ CD4^+^ T cells. (**F**) iSell^Tomato^ mice were infected with AdV and CD4^+^Tomato^+^ T cells were sorted at day 10 post infection. T cells was co-cultured with AdV-GFP infected organoids and imaged with or without anti-IFN-γ blocking antibodies. Normalized infected area over time imaged (left) and accumulated GFP levels (right) in indicated conditions. (**G-H**) WT, iCD4^ΔThPOK^ or iCD4^ΔTCRα^ mice were infected with CR6 (or PBS as control), and infected with AdV 10 days later. AdV stool shedding was measured post AdV-infection. Dashed line represents limit of detection. AdV genome copies per gram of stool of iCD4^ΔThPOK^ (**G**) or iCD4^ΔTCRα^ (**H**) mice relative to WT controls over time. Data are expressed as mean of individual mice, n = 2–9 (minimum two independent experiments in A, C, D, E, F and H. One experiment in B and G). p values as indicated, one-way ANOVA plus Bonferroni test in D, E, G (log10 transformed) and H (log10 transformed). Student t-test in A, B, C, (log10) and F.

To determine if the enhanced IFN-γ production we observed in virus-recruited CD4^+^ T cells (see Figure 3C above) mediates the control of AdV infection, we employed neutralizing anti-IFN-γ antibodies in the co-culture organoid system. CD4^+^Tomato^+^ IE T cells were sorted from AdV-infected iSell^Tomato^ mice 10 days post infection and co-cultured with AdV-GFP infected organoids with or without of anti-IFNγ. Sorted Tomato^+^ T cells inhibited viral replication in the control condition, but not in the presence of neutralizing anti-IFNγ, indicating that IFN-γ derived from virus-recruited CD4^+^ T cells exert an important function in controlling AdV replication (Figure 7F). Conversely, exogenous IFN-γ added to AdV-infected organoids readily suppressed AdV replication (Figure S2D).

Finally, we addressed whether the cross-protective activity observed for CR6-recruited CD4^+^ IE T cells in the AdV-infected organoid system could be recapitulated *in vivo.* Tamoxifen-treated iCD4^ΔThPOK^ or wild-type control mice were infected with CR6 or vehicle, and 10 days later, infected with AdV. WT mice infected with CR6 prior to AdV-infection displayed accelerated AdV clearance compared to vehicle-treated mice, reinforcing the possibility of cross-protection. This phenomenon was dependent on ThPOK expression by CD4^+^ T cells as tamoxifen-treated CR6-infected iCD4^ΔThPOK^ mice did not show reduction in stool AdV titers post infection (Figure 7H). Additionally, as observed during primary AdV infection, initial engagement of the TCR on CD4^+^ T cells during the primary CR6-infection was also necessary for cross-protection against AdV, as CR6-infected iCD4^ΔTCRα^ mice displayed increased AdV titers when compared to wild-type control mice (Figure 7I). Furthermore, excising TCRα from CD4^+^ T cells 6 days post CR6-infection did not impact cross-protection by the recruited CD4^+^ T cells against AdV infection (Figure S2E). Hence, ThPOK-dependent CCR9^+^Ly6A^+^CD4^+^ T cells induced by AdV or CR6 are recruited to the IE compartment, acquiring an increased IFN-γ production that controls AdV viral replication *in vivo* and *in vitro.*

## Discussion

By using a novel mouse genetic strategy allowing the fate-mapping of newly recruited polyclonal T cells to the intestinal tissue, our study uncovered an important role of CD4^+^ IELs in the protection of enteric viral infections. The fate-mapping approach also revealed that while MNV (CR6) and AdV, viruses with tropism for the epithelium, induced a robust CD4^+^ T cell recruitment to the epithelium, CW3 and Reovirus T1L, viruses that predominantly infect cells in the LP, preferentially recruited CD8αβ T cells to both LP and IE compartments. Such intratissue specialization, directly associated with the site of insult, highlights distinct lymphocyte differentiation strategies, as recently demonstrated during steady state (London *et al.*, 2021) and during infection (Kiner et al., 2021).

Upon arriving in the IE compartment at steady state, recruited peripheral CD103^-^ CD4^+^ T cells gradually acquire an IEL phenotype, progressively acquiring CD103 and CD8aa expression (Bilate *et al.*, 2020; London *et al.*, 2021). During this process, a progressive loss of ThPOK allows for increased imprinting by other transcription factors including Runx3 and T-bet, resulting in acquisition of a cytotoxic machinery (London *et al.*, 2021; Mucida *et al.*, 2013; Reis *et al.*, 2014; Reis *et al.*, 2013). As virus-induced cytotoxic CD4^+^ effector T cells have previously been identified in several murine models (Brien et al., 2008; Hou et al., 1992; Stuller et al., 2010) as well as human viral infections (Aslan et al., 2006; Swain et al., 2012; van Leeuwen et al., 2004; Zaunders et al., 2004). IEL differentiation would presumably facilitate response to viruses (Mucida *et al.*, 2013; Reis *et al.*, 2013). While we observed preferential epithelial recruitment of CD4^+^ T cells in the context of AdV or chronic MVN infections, acquisition of an IEL program was prevented and CD4^+^ T cells instead differentiated into IFNγ-producing CCR9^+^Ly6A^+^ cells functionally dependent on continuous ThPOK expression. ThPOK has been previously shown to modulate T_H_1 phenotype during effector differentiation (Vacchio et al., 2014). Additionally, a role for ThPOK in the “functional fitness” of CD4^+^ T cells during responses to systemic LCMV was recently reported (Ciucci et al., 2019), supporting a role for this transcription factor in anti-viral CD4 T cell responses.

In contrast to IEL differentiation pathway (London *et al.*, 2021), we observed that AdV-recruited IE CD4^+^ T cells acquired a mixed T_H_1 and CTL effector phenotype with most of the cells expressing CD69, resembling some characteristics associated to tissue resident memory (T_RM_) though lacking CD103 expression (Mackay et al., 2013; Masopust and Soerens, 2019). Furthermore, a recent study reported that CD103^-^ T_RM_ had increased proliferative potential and enhanced function but could also readily modulate their phenotype upon relocation, in contrast to CD103^+^ T_RM_ (Christo et al., 2021). Runx3, which has been previously associated with CTL (Lotem et al., 2013), IEL (Reis *et al.*, 2013) and T_RM_ (Milner et al., 2017) differentiation, is not required for upregulation of CCR9 or Ly6A by virus-recruited CD4^+^ T cells, but regulates IFN-γ production by these cells. Hence, although the differentiation of IE-recruited CD4^+^ T cells upon virus infection appears to follow a distinct path than T_RM_ and IEL CD4^+^ T cells, it could be related to CTL CD4^+^ T cell responses observed in other tissues, a possibility that requires additional investigation supported by our observations reported here.

Another characteristic of peripheral T cell recruitment to the IE compartment is clonal expansion and progressive loss of TCR diversity as they undergo IEL differentiation during homeostasis (Bilate *et al.*, 2020). In contrast to these steady state observations, virus-recruited IE CD4^+^ T cells displayed a clonally diverse TCR population, which is comparable to what has been described for peripheral CD4^+^ T cells upon LCMV infection (Khatun et al., 2021). Nevertheless, parallel to our observations under steady state (Bilate *et al.*, 2020), while TCR expression was required for initial differentiation and function of IE-recruited CD4^+^ IELs, TCR removal post infection did not impact the capacity of virus-recruited CD4^+^ T cells to control viral replication. This observation, in addition to reinforcing the notion that IE T cells may depend less on TCR engagement to exert their function than peripheral T cells (Bilate *et al.*, 2020), may explain their ability to cross protect against unrelated viruses *in vivo* and in our organoid model. Cross-protection between unrelated antigens have previously been reported regarding tissue-resident CD8αβ^+^ T cells in a skin-infection model of herpes simplex virus (Ariotti et al., 2014).

A relevant point in our studies is that functional IE-recruited CD4^+^ IELs were primarily found in chronic viral model, raising the possibility that targeting strategies to enhance the function of this population may benefit viral control. Primary or secondary (cross-protection) viral clearance data indicated the functional relevance of transcription factors associated with the development of CCR9^+^Ly6A^+^CD4^+^ T cells, or their capacity to secrete IFN-γ, observations related to previous reports describing a crucial role for this cytokine in the human adenovirus replication (Mistchenko et al., 1989). Our study indicates that enteric viral infections promote distinct T cell responses with intra-tissue specialization within the intestine, with previously unappreciated characteristics of TCR repertoire and requirements, surface markers and functional adaptation. Follow up studies should address whether similar pathways exist beyond the intestine and if this information can be translated to human virus infections.

## Materials and methods

### Animals

Animal care and experimentation were consistent with NIH guidelines and were approved by the Institutional Animal Care and Use Committee at the Rockefeller University. *Rosa26^CAG-LSL-tdTomato-WPRE^* (007914), *Zbtb7b*^fl/fl^ (009369), *Cd4*^Cre-ERT2^ (022356) and *Tbx21*^fl/fl^ (022741) mice were purchased from Jackson Laboratories and housed in our facility. Trac^f/f^ mice were kindly provided by A. Rudensky (MSKCC). *Runx3*^fl/fl^ (008773) mice were provided by T. Egawa (Washington University in St. Louis). *Sell*^Cre-ERT2^ mice were provided by M. Nussenzweig (Merkenschlager *et al.*, 2021). Several of these lines were interbred in our facilities to obtain the final strains described elsewhere in the text. Genotyping was performed according to the protocols established for the respective strains by Jackson Laboratories or by donor investigators. Mice were maintained at the Rockefeller University animal facility under specific pathogen-free (SPF) conditions. Both male and female littermates were used, age 7-12 weeks old.

### Tamoxifen Treatment

Tamoxifen (Sigma) was dissolved in corn oil (Sigma) and 10% ethanol, shaking at 37°C for 30 min-1 h. Two doses of Tamoxifen (5 mg/dose) were administered to mice via oral gavage at 50 mg/mL, 3 days and 1 day before viral infection.

### Oral infection with *Listeria monocytogenes*

*L. monocytogenes* 10403S-inlA strain expressing full-length OVA (Lm-OVA) were grown overnight in brain heart infusion media. Lm-OVA was provided by L. Lefrançois. Mice were infected with 10^9^ colony forming units (CFU) 24h after oral treatment with 20 mg of Streptomycin (Sigma-Aldrich) diluted in water. At day 9 post-infection, intraepithelial lymphocytes were harvested and stained with LLO-tetramer (NIH) and analyzed by flow cytometry.

### Oral infection with viruses

WT mouse adenovirus-2(AdV) and AdV-GFP [previously called “MAdV-2.IXeGFP (Wilson *et al.*, 2017)] were propagated in the mouse rectal carcinoma CMT-93 cell line(ATCC CCL-223), purified, and quantified as previously described (Gounder et al., 2016). Mice were infected with 10^7^ infectious units of WT AdV in 100 ul PBS by oral gavage. Reovirus T1L (T1L) was provided by T. Dermody (University of Pittsburgh) and propagated in L929 cell line, purified and quantified as previously described (Kobayashi et al., 2010). T1L. Mice were infected with 10^8^ plaque forming units (pfu) of T1L in 100 ul PBS by oral gavage. Murine norovirus (MVN) CW3 and CR6 were provided by K. Cadwell (New York University) and was propagated in RAW264.7 cell line, purified and quantified as previously described (Kernbauer *et al.*, 2014). Mice were infected with 3×10^6^ pfu of MNV CW3 or CR6 in 100 ul PBS by oral gavage.

### Adenovirus fecal shedding

1 or 2 stool pellets were collected at the days post infection as indicated in the figure legends and frozen at −80°C. On the day of processing, stool samples were weighed and processed with QIAamp Fast DNA Stool Mini Kit (Qiagen) for DNA extraction following the manufacturer’s instructions. Samples were measured for DNA content by NanoDrop (Thermo Scientific). RT-qPCR was performed using PowerSYBR Green (Applied Biosystems) with AdV specific primers; FW: 5’-GTCCGATTCGGTACTACGGT-3’; RV: 5’-GTCAGACAACTTCCCAGGGT-3’, at an annealing temperate of 55°C and for 40 cycles on a QuantStudio 3 RT PCR System (Applied Biosystems). Genomic copies were determined by correlation to an AdV DNA standard and normalized to DNA input and stool weight. Stool from uninfected mice was used as a negative control.

### Isolation of intestinal T cells

Intraepithelial and lamina propria lymphocytes were isolated as previously described (Reis *et al.*, 2013). Briefly, small intestines were harvested and washed in PBS and 1 mM dithiothreitol (DTT) followed by 30 mM EDTA. Intraepithelial cells were recovered from the supernatant of DTT and EDTA washes. Lymphocytes from lamina propria were obtained after collagenase digestion of the tissue. Mononuclear cells were isolated by gradient centrifugation using Percoll. Single-cell suspensions were then stained with fluorescently labeled antibodies for 25 min at 4° C prior to downstream flow cytometry (analysis or sorting) as specified in figure legends.

### Staining Strategy

The following gating strategy was utilized to examine CD4^+^ T cells: single live lymphocytes (based on size and live/dead fixable dye Aqua stain), CD45^+^, TCRγδ^-^, TCRβ^+^, CD8β^low/–^, CD4^+^. For analysis of iSell^Tomato^, the following gating strategy was added: CD62L^-^ and Tomato^+^. For single-cell sorting of cells subjected to scTCRseq the following gating strategy was used: single live lymphocytes, CD45+, TCRγδ^-^, TCRβ^+^, CD4^+^, CD62L^-^, Tomato^+^, Ly6a^+/–^, CD103^+/–^, CD8α^+/–^. For sorting of cells subjected to bulk RNaseq we used single live lymphocytes CD45^+^, TCRγδ^-^, TCRβ^+^, CD8β^-^, CD4^+^, Tomato^+^ and CD62L^-^. For sorting of T cells subjected to organoid co-cultures, we gated on single live lymphocytes CD45^+^ TCRγδ^-^, CD8β^-^, CD62L^-^, MHCII^-^, CD11b^-^, CD11c^-^, CD19^-^, NK1.1^-^, Tomato^+^ and CD5^+^.

### Intracellular Staining and Flow Cytometry

For analysis of cytokine secretion, total mononuclear cells isolated from the epithelium were plated in 48-well plates and incubated at 37° C with 100ng/mL phorbol 12-myristate 13-acetate (PMA, Sigma), 200ng/mL ionomycin (Sigma) and 2mM monensin (BD Biosciences) for 4 h. Intracellular staining for IFN-γ and granzyme B was conducted in Perm/Wash buffer after fixation and permeabilization in Fix/Perm buffer (BD Biosciences, USA) according to kit instructions. Flow cytometry data were acquired on an LSR-II flow cytometer (Becton Dickinson,USA) and analyzed using FlowJo 10 software package (Tri-Star, USA).

### Organoids and co-cultures

Organoids were established from crypts isolated from adult mouse small intestine and maintained as described previously (Rogoz et al., 2015; Sato *et al.*, 2009). For infection and co-cultures, organoids were removed from the Matrigel by gently pipet off the top media layer and 500 μl of cold culture media containing 10% FBS (Sigma F0926) was added to each well to dissolve the Matrigel. Organoids was washed with cold culture media containing 10% FBS and 30-40 organoids were infected with 10^4^ i.u. of AdV-GFP in 500 μl cold PBS for 20 min on ice. Inoculum was discarded and infected organoids was washed with 15 ml of T-cell culture medium (TCM) [RPMI 1640, 10% FBS, 1% Pen/Strep (Gibco), 1% L-glutamine (Gibco), 1% Sodium Pyruvate (Gibco), 2% Non-essential Amino Acids (Gibco), 2.5% 1M HEPES (Gibco), 50μM 2-Mercaptoethanol (Sigma)]. Then, working on ice, 30 μl of 2-3×10^4^ sorted CD4^+^Tomato^+^ T cells in TCM was carefully added to 30 ul of 8-12 infected organoids in 30 μl TCM, next we added 40 ul of ice cold Matrigel. 100 μl of the T cell/organoid/Matrigel mixture was immediately added to a preheated (37°C) 96-well culture plate with glass bottoms (MatTek). The culture plate was incubated for 20-30 min in 37°C to let the Matrigel polymerize. Next, we added 200 μl of TCM containing 50 ng/ml recombinant murine EGF, 100 ng/ml recombinant murine Noggin, 500 ng/ml recombinant human R-spondin, 10 ng/ml recombinant murine IL-2 (R&D), 5 ng/ml recombinant murine IL-7 (R&D) and 10 ng/ml recombinant murine IL-15/IL-15R (R&D) on top of the polymerized Matrigel. For IFN-γ neutralizing experiments, we also added 10 ng/ml of anti-IFN-γ (BD Biosciences).

### Organoid analysis and T cell tracking

Live imaging was performed on the CellVoyager (Yokagawa/Olympus) spinning disk confocal microscope at 37°C and with 5% CO2. Co-cultures were imaged in 96-well plates with glass bottoms (MatTek). 10 to 12 z-stacks of 1.52 μm step-size were acquired every 6-7 min for 48 hours. GFP signal above background per organoid was used to measure the GFP expression area for each z-stack and time point. Organoid area was determined with brightfield images for each z-stack and time point and used to normalize the GFP expression area for each respective z-stack and time point. For T cell tracking, hyperstacks were made of each z-stack, time point and channel. T cell (Tomato^+^) and infected epithelial (GFP^+^) tracking was obtained with TrackMate 6.0.2 (Tinevez et al., 2017) on hyperstacked images and over time, cell diameter was set at 14 μm to obtain optimal tracking. Tomato^+^ T cell tracking 3D coordinates was analyzed together with GFP^+^ infected epithelial 3D coordinates. If a Tomato^+^ track point was within 20μm radius of a GFP^+^ track point in any dimension at a specific given time, that track point was considered as an interacting point. A minimum of 8 sequential interacting points in a specific T cell track were used to calculate the cell average velocity. Reconstruction of images were done using a Python script and the skimage v.0.19 package. Image quantifications and modulations were made in ImageJ 2.1 and all image calculations were made in R studio 1.2.5.

### Single-Cell TCR Sequencing

Single cells were index-sorted using a FACS Aria into 96-well plates containing 5μL of lysis buffer (TCL buffer, QIAGEN 1031576) supplemented with 1% β-mercaptoethanol) and frozen at −80° C prior to RT-PCR. RNA and RT-PCRs for TCRβ were prepared as previously described (Dash et al., 2011). PCR products for TCRβ were multiplexed with barcodes and submitted for MiSeq sequencing (Han et al., 2014) using True Seq Nano kit (Illumina). Fastq files were de-multiplexed and paired-end reads were assembled at their overlapping region using the PANDASEQ (Masella et al., 2012) and FASTAX toolkit. Demultiplexed and collapsed reads were assigned to wells according to barcodes. Fasta files from MiSeq sequences were then aligned and analyzed on IMGT (http://imgt.org/HighV-QUEST) (Brochet et al., 2008). Cells with identical TCRβ CDR3 nucleotide sequences were considered as the same clones.

### Bulk RNA-seq Library Preparation

Sorted cells (800 cells) were lysed in a guanidine thiocyanate buffer (TCL buffer, QIAGEN) supplemented with 1% β-mercaptoethanol. RNA was isolated by solid-phase reversible immobilization bead cleanup using RNAClean XP beads (Agentcourt, A63987), reversibly transcribed, and amplified as described (Trombetta et al., 2014). Uniquely barcoded libraries were prepared using the Nextera XT kit (Illumina) following the manufacturer’s instructions. Sequencing was performed on an Illumina NextSeq500 for a total yield of 400M reads.

### Bulk RNA-seq Analysis

Raw fastq files were processed by using the mouse transcriptome (gencode M23) with the kallisto (v0.46) software (Bray et al., 2016). Analysis of transcript quantification was performed at the gene level by using the sleuth (v0.30) package for R (Pimentel et al., 2017). In short, we modeled batch effect and our experimental design using the sleuth_fit function and detected differentially expressed genes between all groups by the likelihood ratio test (LRT). To determine significantly expressed genes between group pairs, we used the wald-test function. All downstream analysis was made using genes below adjusted p value of 0.05. TCRα and TCRβ CDR3 sequences were reconstructed *in silico* using MixCR software (Bolotin et al., 2015) and the extracted sequences were analyzed with the Immunarch package (v0.6.5) for R (10.5281/zenodo.3893991).

### Statistical Analyses

Statistical analysis was carried out using GraphPad Prism v.9. Flow cytometry analysis was carried out using FlowJo software. Data in graphs show mean ± SEM and p values < 0.05 were considered significant. Repertoire diversity was analyzed by the Diversity 50 (D50). Diversity 50 (D50) was calculated using Excel version 16 as the fraction of dominant clones that account for the cumulative 50% of the TCRβ CDR3s identified. GraphPadPrism v.9 was used for graphs and Adobe Illustrator 2020 used to assemble and edit figures.

## Acknowledgments

We are grateful to A. Rogoz and S. Gonzalez for exceptional animal care, mouse colony management and genotyping; Yasmeen Khan, the Genomics Core, and additional Rockefeller University employees for continuous assistance. We thank the NIH tetramer facility for providing the LLO tetramer. We thank A. Bilate for providing advice on the TCR sequencing and editing the manuscript, A. Lockhart for editing the manuscript and all the members of the Mucida, Victora (Rockefeller) and Lafaille (NYU) labs for fruitful discussions.

This work was supported by the Black Family Metastasis Center, the Burroughs Wellcome Fund PATH Award, the Mathers Foundation, the Pershing Square Foundation, the FASI/FARE Consortium and National Institute of Health grants R21AI144827, R01DK113375 and R01DK093674 (D.M.). R.P. was supported by the Swedish Research Council and The Sweden-America Foundation.

## Author contributions

R.P. designed and performed experiments and wrote the manuscript. R.P. and T.B.R.C. performed all bioinformatics analyses and T.B.R.C. assisted with interpretation of sequencing data. R.P. and M.L. performed sample and library preparation of bulk RNA-seq. R.P., M.L. and B.R. performed single-cell PCRs and multiplexing of samples for single-cell TCR-seq. M.L. and T.B.R.C. helped with analysis of single cell TCR-seq by MiSeq. R.P. and J.B.A. setup and performed organoid experiments. J.G.S. purified and provided WT and engineered adenovirus. D.M. conceived and supervised the research and wrote the manuscript. All authors edited the manuscript.

---

**Supplementary figure 1. AdV infection and T cell dynamics post viral infections.** (**A**) Mice were infected with 10^7^ i.u. of AdV-mCherry and ileum were analyzed at day 6 post infection. Scale bar represents 60 μm. (**B**) Left: iSell^Tomato^ mice were treated with tamoxifen and mesenteric lymph nodes (mLN) were analyzed 3 days later. Right: tamoxifen-treated iSell^Tomato^ mice were orally infected with 10^9^ cfu of Listeria monocytogenes and lymphocytes were isolated from the epithelium (IE) and the frequency of Listeriolysin O (LLO)-tetramer^+^ Tomato^+^ cells among CD4^+^ T cells was analyzed 9 days post-infection (**C**) iSell^Tomato^ mice were orally infected with 10^7^ i.u. of AdV, 10^8^ pfu of T1L, 3×10^6^ pfu of CR6 or 3×10^6^ of CW3, TCRβ^+^CD4^+^CD62L^-^ and TCRβ^+^CD8αβ^+^CD62L^-^ LP cells were analyzed for tomato expression 10 days post infection. (**D**) iSell^Tomato^ mice were orally infected with 10^8^ pfu of T1L, 3×10^6^ pfu of CR6 or 3×10^6^ of CW3, TCRb^+^CD4^+^CD62L^-^Tomato^+^ IELs were analyzed for CD69, CD103, CD244 and CD8αα expression 10 days post infection. Data are expressed as mean of individual mice (n = 3–5 of two independent experiments). p values as indicated, One-way ANOVA plus Bonferroni test in C.

**Supplementary figure 2. Analysis of iSell^Tomato^ mice 30 days post AdV infection and the role of CCR9^+^Ly6A^+^ T cells, IFNγ and TCRα in AdV-recruited T cells.** (**A**) iSell^Tomato^ mice were orally infected with 10^7^ i.u. of AdV and TCRβ^+^CD4^+^CD62L^-^ IELs were analyzed for Tomato, CD8αα and CD103 expression 30 days post infection. (**B**) Viral genome copies of AdV per gram of feces of AdV-infection over time of WT and iCD4^ΔTCRα^ mice treated with tamoxifen at day 6 and 8 post infection. (**C**) Viral genome copies of AdV per gram of feces of AdV-infection of WT or iCD4^ΔTCRα^ mice after primary infection (day 10 post infection) or secondary infection (60 days post primary infection, mice were treated with tamoxifen 1 and 3 days before secondary infection). (**D**) 8-10 organoids were infected with 10^4^ i.u. of AdV-GFP and treated with or without 10 ng/ml of IFNγ. Organoids were homogenized into a single cell suspension with TrypLE and stained for EpCAM. Cells were analyzed by flow cytometer. (**E**) Viral genome copies of AdV per gram of feces of AdV-infection over time of WT and iCD4^ΔTCRα^ mice treated with tamoxifen at day 6 and 8 post MNV infection. Data are expressed as mean of individual mice (n = 3–5 of two independent experiments in A. Pooled data from two independent experiments in B and data from one experiment in E). Data are expressed of one experiment with 8-10 organoids in D. p values as indicated, Student’s t-test.

**Supplementary movie 1.** Small intestine organoids were infected with 10^4^ i.u. of AdV-GFP and imaged for 48 hours.

**Supplementary movie 2.** Small intestine organoids were infected with 10^4^ i.u. of AdV-GFP and co-cultured with CD4^+^Tomato^+^ IELs from AdV-infected mice. Co-cultures was imaged for 50 hours.

**Supplementary movie 3.** Small intestine organoids were infected with 10^4^ i.u. of AdV-GFP and co-cultured with CD4^+^Tomato^+^ IELs from CR6-infected mice. Co-cultures was imaged for 50 hours.

**Supplementary movie 4.** Small intestine organoids were infected with 10^4^ i.u. of AdV-GFP and co-cultured with CD4^+^Tomato^+^ IELs from T1L-infected mice. Co-cultures was imaged for 49 hours.

