## Supplementary figures and images for "Newly recruited intraepithelial Ly6A^+^CCR9^+^CD4^+^ T cells protect against enteric viral infection"

### Supplementary Figure 1

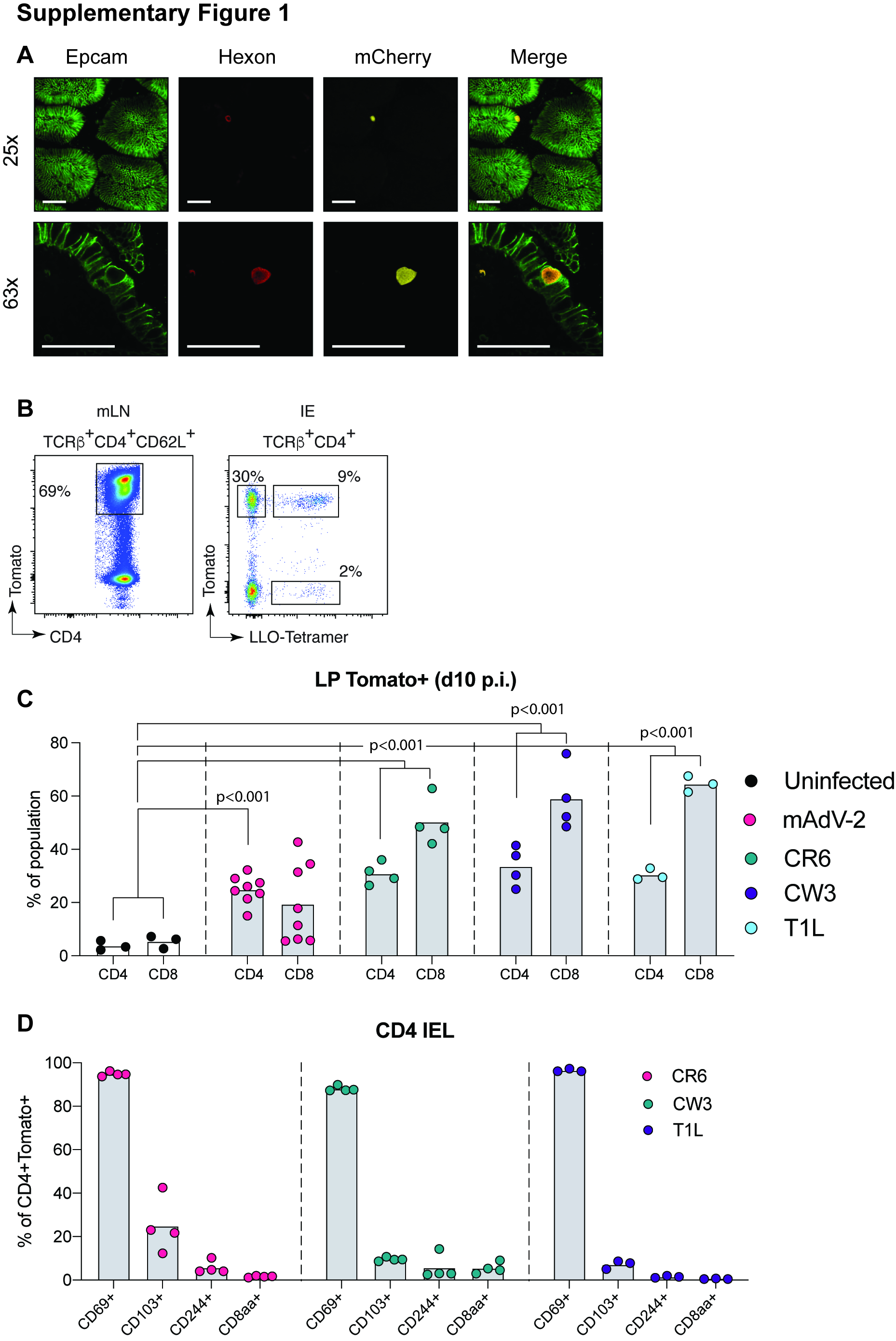

### Supplementary Figure 2

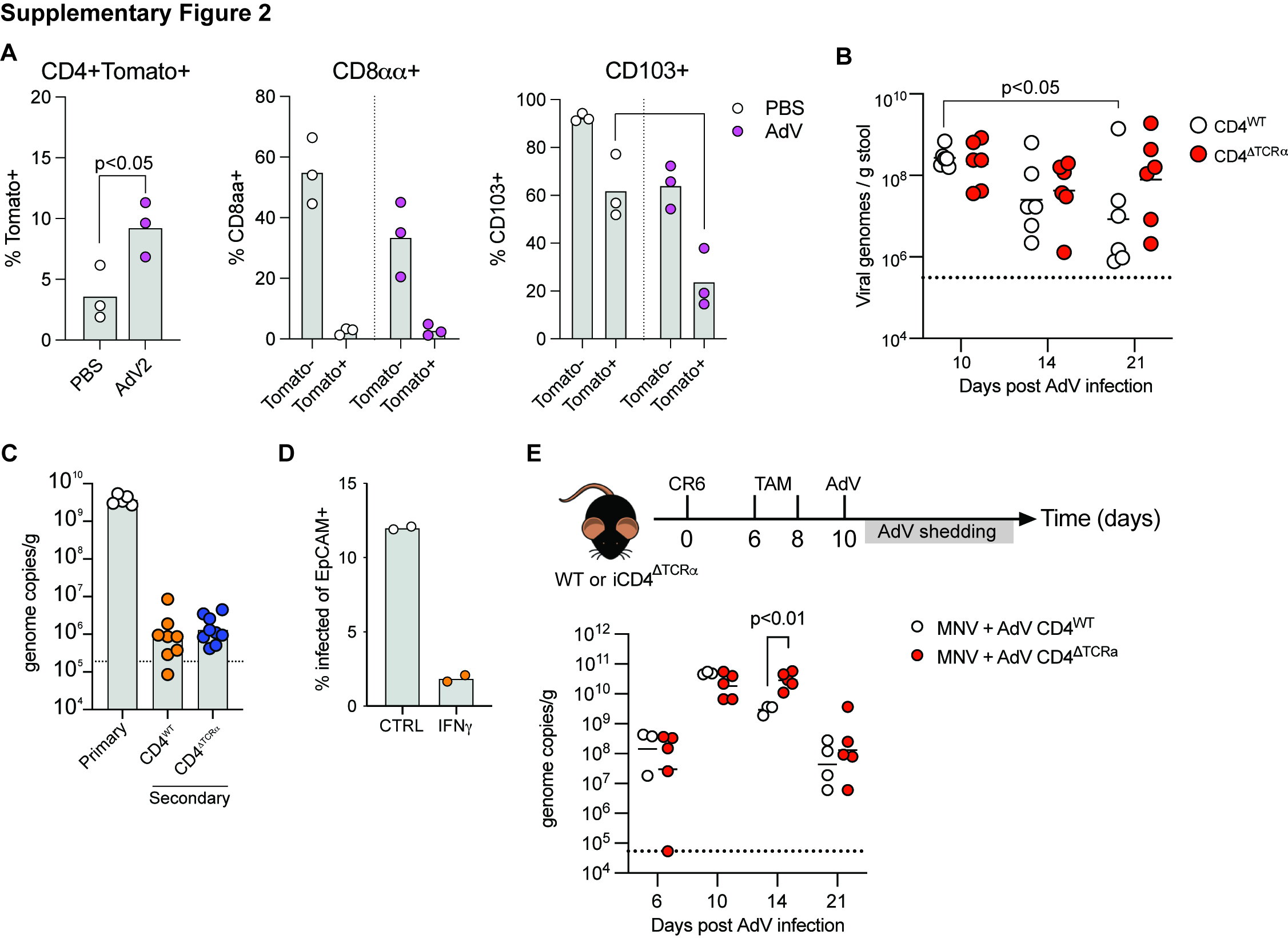
